## Supplementary Figures for "Multimodal monitoring of human cortical organoids implanted in mice using transparent graphene microelectrodes reveal functional connection between organoid and mouse visual cortex"

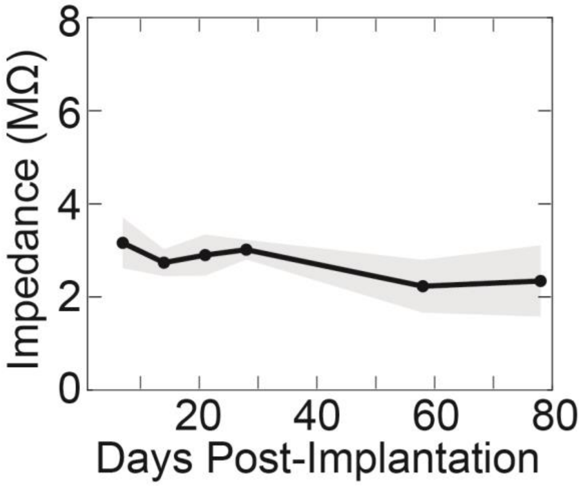


**Supplementary Figure 1.** Graphene microelectrode impedance over time for a representative animal. Results are shown as mean ± *sdv* across 16 channels.

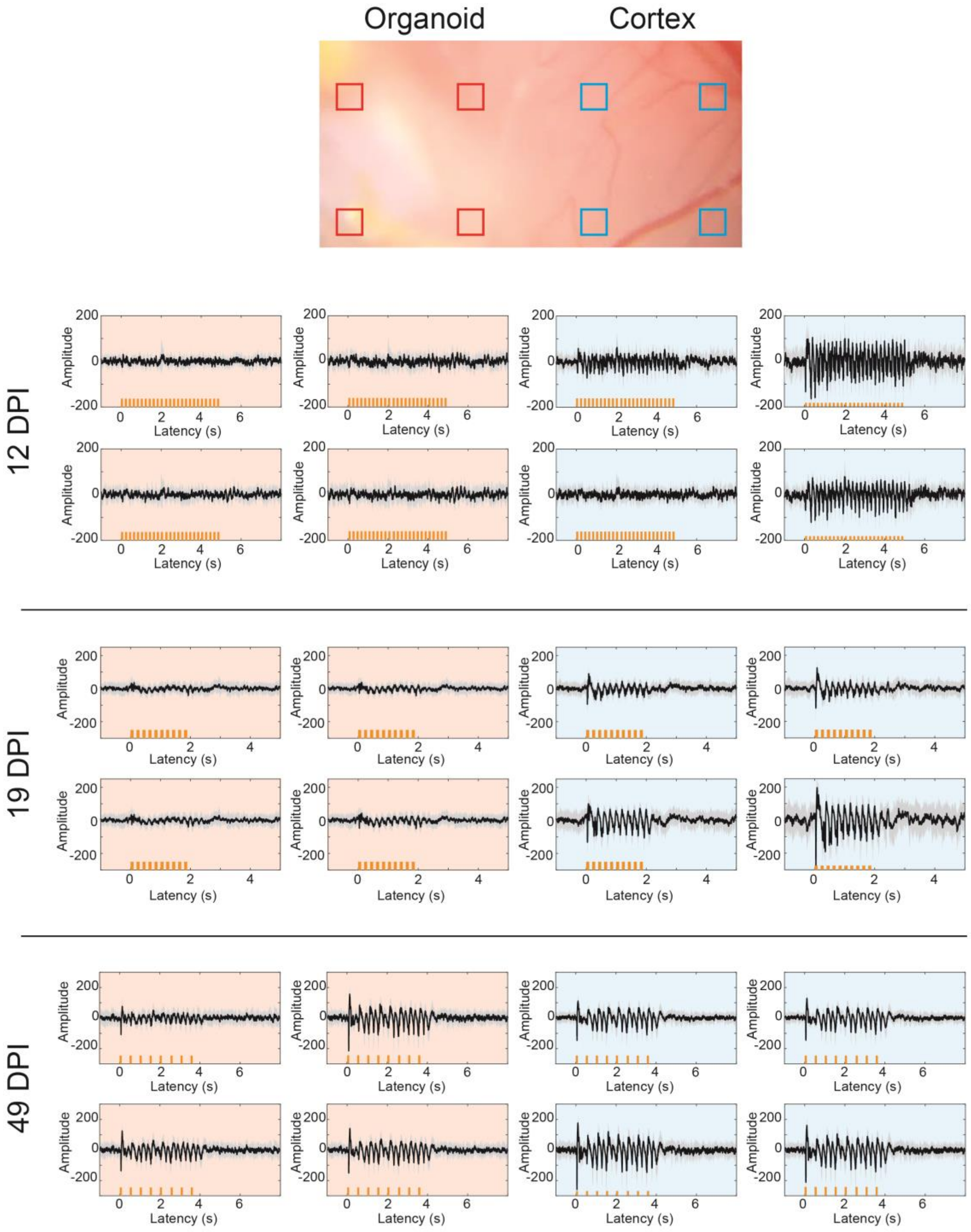


**Supplementary Figure 2.** Increase of local field potential (LFP) amplitude over time. LFP are shown as mean ± *sdv* (n=10 trials for 12 and 19 dpi and n=20 trials for 49 dpi). The brightfield image shows the relative location of the channels shown; channels covering the organoid implantation area are outlined in red.

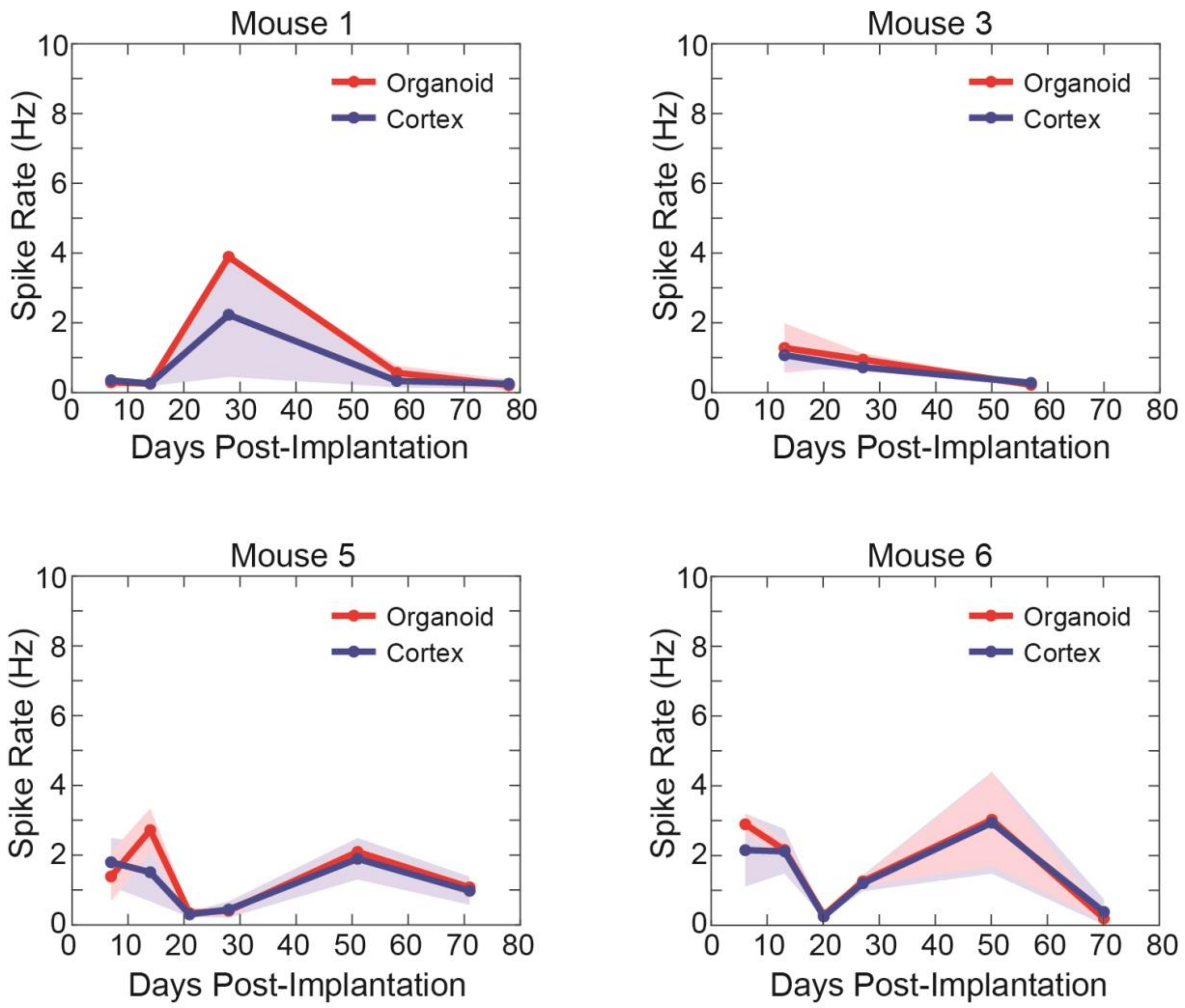


**Supplementary Figure 3**. Change in MUA spike rates over time for four mice. Rates are shown as mean ± *sdv* (of 5±3 channels for mouse 1 and 3 and 13±3 channels for mice 5 and 6) for channels overlaying cortex or organoid implantation. A MUA event threshold of -4**sdv* was used for all recordings.

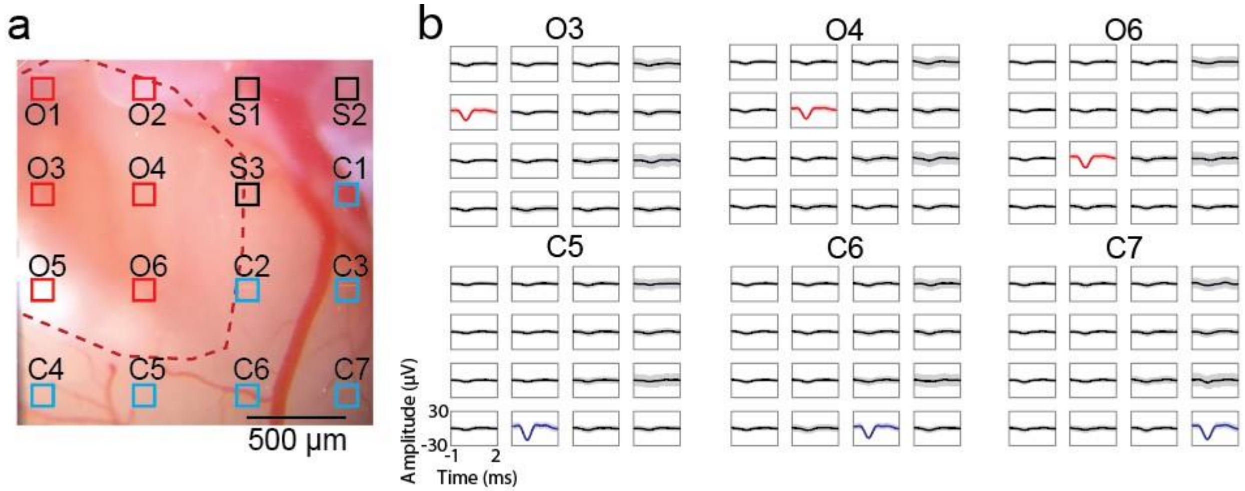


c

| Ch | S1 | S3 | O2 | O1 | O3 | O4 | O5 | C4 | C5 | O6 | C6 | C7 | C3 | C2 | C1 | S2 |
|---|---|---|---|---|---|---|---|---|---|---|---|---|---|---|---|---|
| S1 | 147 | 1 | 2 | 2 | 0 | 1 | 2 | 1 | 1 | 0 | 1 | 0 | 0 | 0 | 0 | 0 |
| S3 | 1 | 80 | 2 | 0 | 1 | 2 | 1 | 0 | 1 | 1 | 0 | 0 | 0 | 0 | 0 | 0 |
| O2 | 2 | 2 | 118 | 0 | 1 | 1 | 2 | 0 | 1 | 0 | 1 | 0 | 0 | 0 | 0 | 0 |
| O1 | 2 | 0 | 0 | 97 | 0 | 0 | 2 | 0 | 0 | 0 | 0 | 0 | 0 | 0 | 0 | 0 |
| O3 | 0 | 1 | 1 | 0 | 127 | 1 | 1 | 1 | 0 | 0 | 0 | 0 | 0 | 0 | 0 | 0 |
| O4 | 1 | 2 | 1 | 0 | 1 | 213 | 4 | 0 | 1 | 1 | 1 | 0 | 0 | 0 | 0 | 0 |
| O5 | 2 | 1 | 2 | 2 | 1 | 4 | 139 | 0 | 1 | 0 | 1 | 3 | 0 | 0 | 0 | 0 |
| C4 | 1 | 0 | 0 | 0 | 1 | 0 | 0 | 69 | 0 | 0 | 1 | 0 | 0 | 0 | 0 | 0 |
| C5 | 1 | 1 | 1 | 0 | 0 | 1 | 1 | 0 | 55 | 0 | 1 | 0 | 0 | 0 | 0 | 0 |
| O6 | 0 | 1 | 0 | 0 | 0 | 1 | 0 | 0 | 0 | 126 | 0 | 0 | 0 | 0 | 0 | 0 |
| C6 | 1 | 0 | 1 | 0 | 0 | 1 | 1 | 1 | 1 | 0 | 97 | 0 | 0 | 0 | 0 | 0 |
| C7 | 0 | 0 | 0 | 0 | 0 | 0 | 3 | 0 | 0 | 0 | 0 | 36 | 0 | 0 | 0 | 0 |
| C3 | 0 | 0 | 0 | 0 | 0 | 0 | 0 | 0 | 0 | 0 | 0 | 0 | 4 | 0 | 0 | 0 |
| C2 | 0 | 0 | 0 | 0 | 0 | 0 | 0 | 0 | 0 | 0 | 0 | 0 | 0 | 26 | 0 | 0 |
| C1 | 0 | 0 | 0 | 0 | 0 | 0 | 0 | 0 | 0 | 0 | 0 | 0 | 0 | 0 | 30 | 0 |
| S2 | 0 | 0 | 0 | 0 | 0 | 0 | 0 | 0 | 0 | 0 | 0 | 0 | 0 | 0 | 0 | 13 |

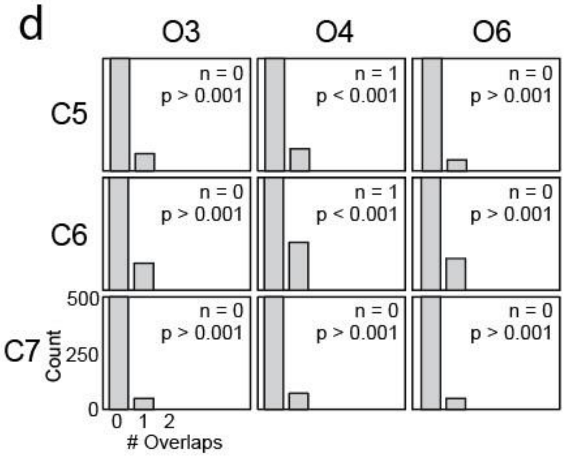


**Supplementary Figure 4.** MUA analysis in three mice to investigate the overlap of signal across channels. (**a**) Brightfield image of mouse cortex with organoid region outlined in red. Red channels are those overlapping the organoid. (**b**) Event-averaged MUA traces show that the MUA events are localized spatially (mean ± *sdv*). (**c**) Table showing the number of overlapping events after binning events into 1 ms windows for a ~100 s spontaneous recording trial. Diagonal shows the number of events detected per channel. Red color channels are channels overlaying the organoid, blue color channels are those overlaying cortex. (**d**) Histograms of the number of overlaps across an organoid channel and cortex channel (shown in plot titles) after circularly shuffling the MUA event trains 10,000 times. P-values were determined by integrating the shuffled counts from the overlap count of the non-shuffled case (n) in panel c to infinity. Large p-values indicate no significant overlap across channels, supporting that the MUA data is independent across channels.

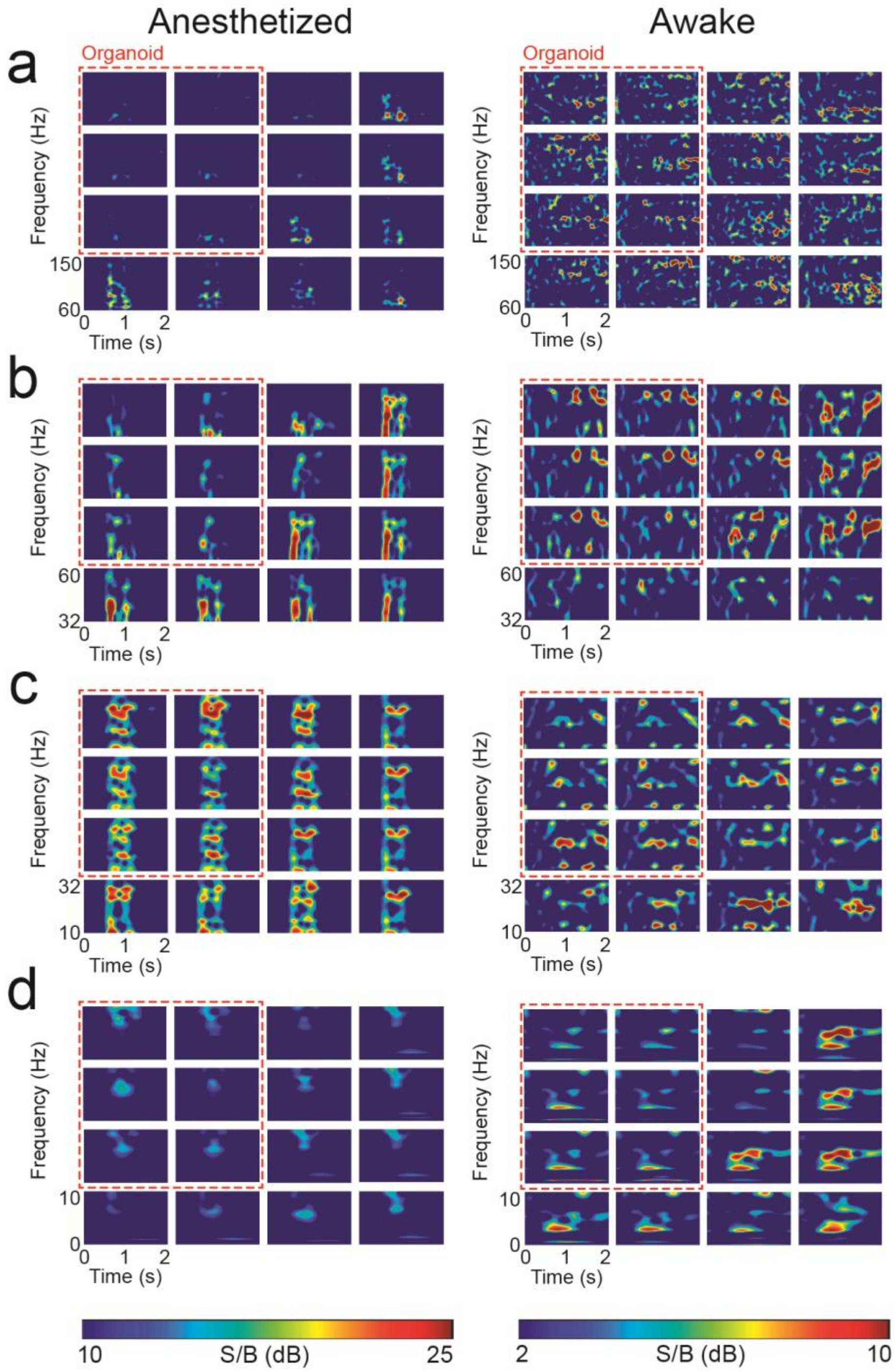


**Supplementary Figure 5.** Spectrograms during representative epochs (same epochs as figure 2c and 2e) for different frequency bands broken into sub-frequency ranges for easier visualization: (**a**) 60-150 Hz, (**b**) 32-60 Hz, (**c**) 10-32 Hz, and (**d**) 0-10 Hz. The red dashed box in all panels delineates channels overlaying the organoid. A discrepancy appears along the organoid border for low and high (> 32 Hz) gamma bands while the mouse was under anesthesia with 1.5% isoflurane.

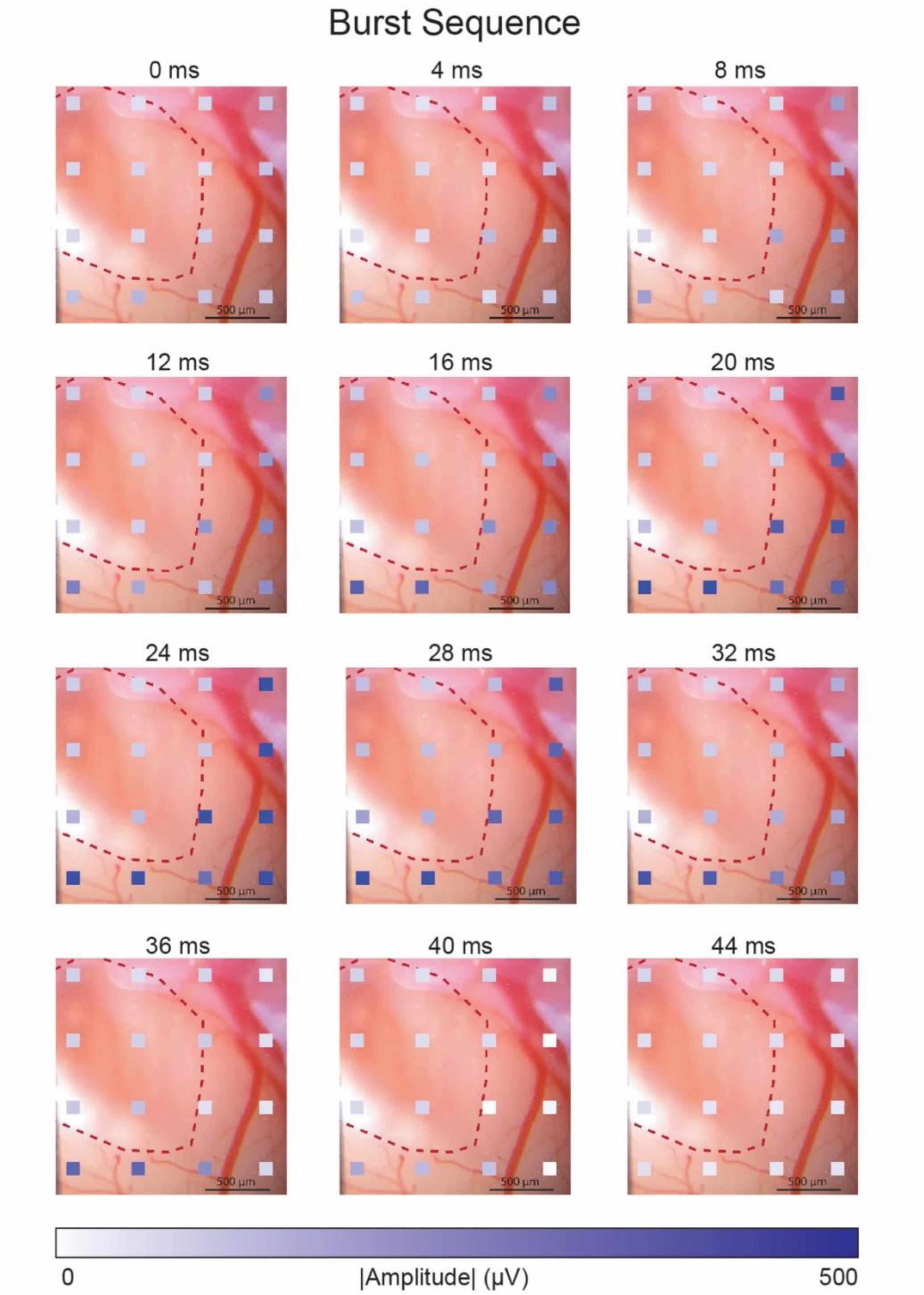


**Supplementary Figure 6.** Local field potential amplitude for all 16 channels during a burst event while the mouse was under anesthesia with 1.5% isoflurane.

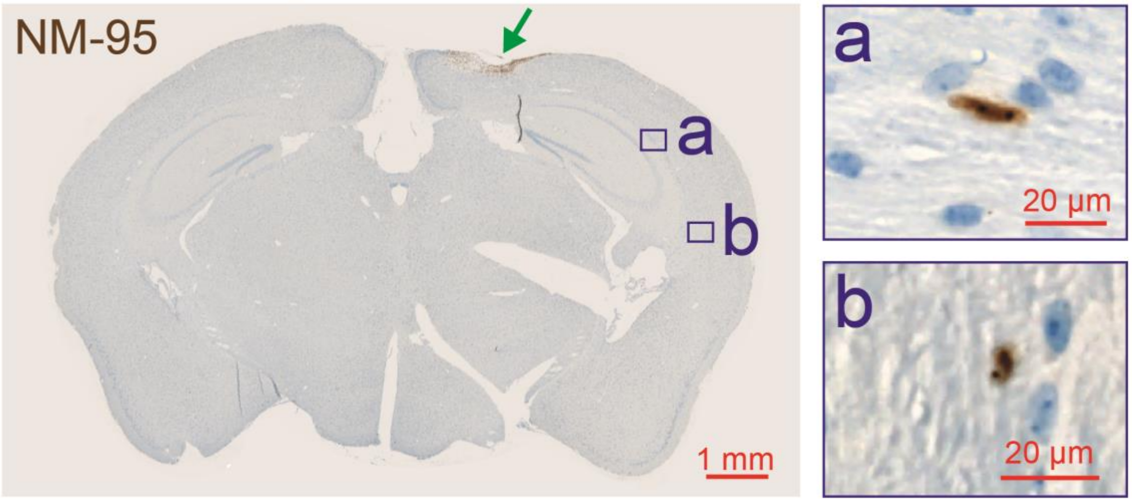


**Supplementary Figure 7**. Human (NM-95-positive) cells were observed at the implantation site (green arrow) and further away from the implantation site (**a, b**); we detected individual human cells along corpus callosum up to ~4 mm away from the implantation site (**b**).

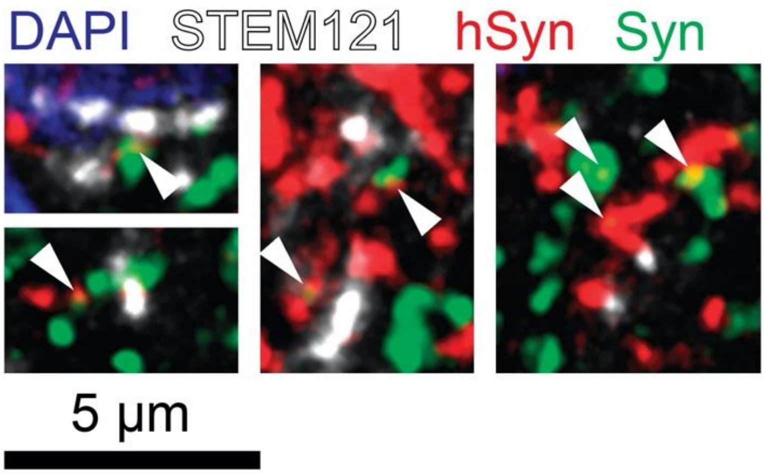


**Supplementary Figure 8.** Puncta that labeled positive for both Syn (green) and hSyn (red) were counted as presynaptic puncta of human origin (yellow, arrowheads). We observed a presence of STEM121 (white) and human presynaptic puncta (yellow, arrowheads) within regions of visual cortex, supporting the organoid extended axonal connections towards and into mouse visual cortex.
