## Supplementary Video 1 Description for "Multimodal monitoring of human cortical organoids implanted in mice using transparent graphene microelectrodes reveal functional connection between organoid and mouse visual cortex"

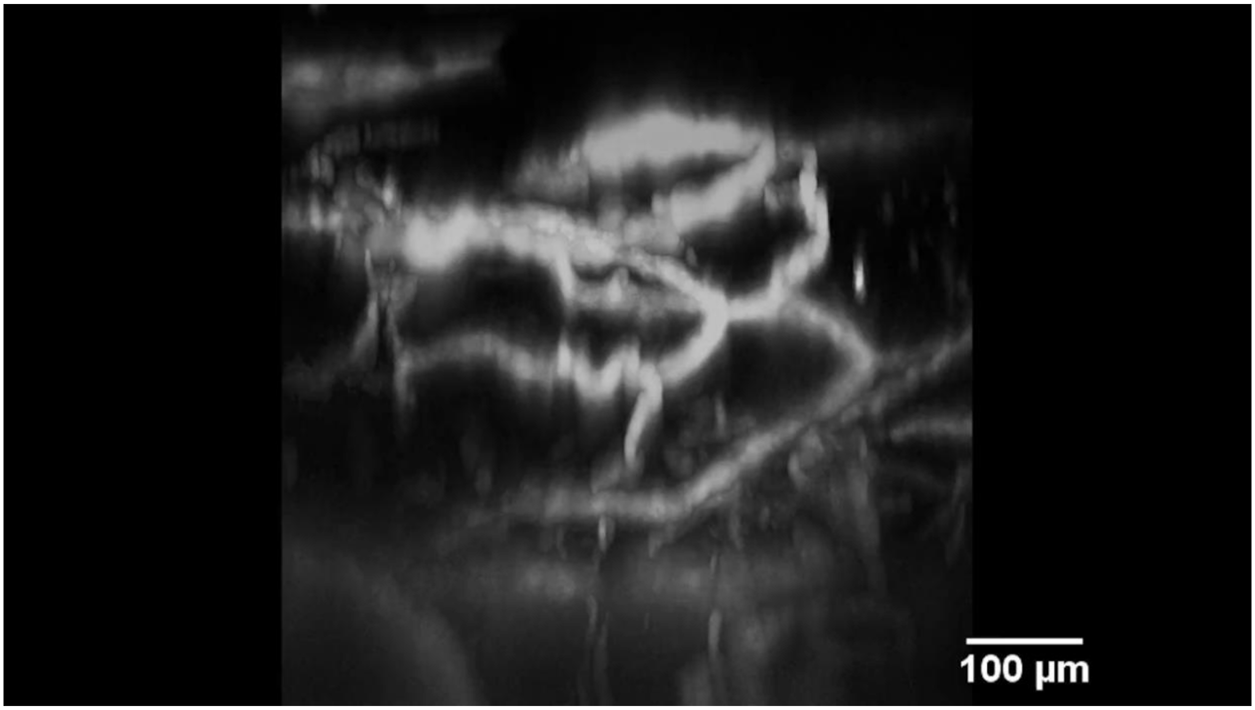


**Supplementary Video 1 (screenshot).** Three-dimensional projection of the vasculature within the organoid implant region. Image was acquired using two-photon microscopy after injection of Alexa Fluor 680 Dextran.
